# *Arabidopsis TITAN LIKE* is required for U12-type intron splicing, especially of AT–AC subtypes

**DOI:** 10.1101/2025.02.25.639993

**Authors:** Tomoko Niwa, Junshin Miyamoto, Nao Iwase, Haruka Iwai, Takaaki Kojima, Takamasa Suzuki

## Abstract

Many eukaryotes possess two types of spliceosomes: the U2-dependent and U12-dependent spliceosomes. The U2-dependent spliceosome processes more than 99% of all introns, whereas the U12-dependent spliceosome acts on only about 0.3% of introns, one-third of which start with AT and end with AC, with the remainder having GT–AG termini. How the U12-dependent spliceosome splices two types of introns with different terminal sequences remains poorly understood. Human CENATAC is a subunit of the U12-dependent spliceosome that is particularly required for the splicing of the AT–AC subtype. The *Arabidopsis* genome contains a single homolog, *TITAN LIKE* (*TTL*), but its function in splicing remains unknown. Here, we generated TTL mutants and isolated two viable alleles, of which we analyzed one, designated *ttl-142*, to investigate TTL function in splicing. *ttl-142* carries a 42-nucleotide deletion that removes 14 amino acid residues from the predicted protein, and homozygous mutants exhibit morphological abnormalities. Most U12-dependent introns were less efficiently spliced in *ttl-142* than in the wild-type, with the splicing of AT–AC introns particularly suppressed. Splicing suppression in *ttl-142* was more extensive than that in a *DROL1* mutant, which carries a mutation in a gene specifically required for AT–AC intron splicing. Conversely, fewer genes showed altered expression levels in *ttl-142* than in *drol1*, and most differentially expressed genes differed between the two mutants. These results suggest that the phenotypes of *ttl-142* and *drol1* mutants may reflect the impairment of distinct spliceosomal functions.

## Introduction

Many eukaryotes, including plants and vertebrates, possess two types of spliceosomes (Turunen et al. 2013). The first is the U2-dependent (major) spliceosome, which comprises five small nuclear RNAs (snRNAs)—U1, U2, U4, U5, and U6—along with numerous associated proteins (Wilkinson et al. 2019). The U2-dependent spliceosome is responsible for splicing nearly 99% of introns. The second type, the U12-dependent (minor) spliceosome, consists of U11, U12, U4atac, U5, and U6atac snRNAs, together with both unique proteins and several shared with the U2-dependent spliceosome (Turunen et al. 2013; Ding et al. 2023). The introns spliced by each spliceosome differ in their consensus sequences at the 5′ splice site and branch point sequences and have long been considered mutually exclusive. Based on these sequence features, approximately 400 introns of the more than 125,000 introns in the Arabidopsis genome have been predicted to be U12-type introns (Szcześniak et al. 2013). Although the GT–AG intron termini are highly conserved among eukaryotes, approximately one-third of U12-type introns spliced by the U12-dependent spliceosome possess AT–AC termini. How the U12-dependent spliceosome processes introns with these distinct boundaries remains unclear. Recently, two protein subunits specifically required for the splicing of AT–AC introns—DROL1 and CENATAC—have been identified (Suzuki et al. 2022; de Wolf et al. 2021).

While screening for *Arabidopsis thaliana* mutants defective in the repression of seed-maturation genes, we identified the *defective repression of the OLE3:LUC 1* (*drol1*) mutant, which carried a mutation in a splicing factor (Suzuki et al. 2018). The *A. thaliana* DROL1 protein is homologous to yeast DIB1 and human Dim1, both subunits of the U5 snRNP, but belongs to a distinct subfamily (Suzuki et al. 2022). Furthermore, comprehensive transcriptomic analysis of the *drol1* mutant revealed that splicing of AT–AC introns was specifically suppressed, suggesting that DROL1 functions as a splicing factor specific to AT–AC introns.

The Dim1 protein, together with PRP8, forms a binding pocket for 5′ splice sites in the B complex of the human U2-dependent spliceosome (Bertram et al. 2017; Zhan et al. 2018). In this complex, the 5′-terminal guanine stacks with a phenylalanine residue in PRP8, while the adjacent uracil is positioned close to Dim1 (Bertram et al. 2017). Structural analyses have suggested that Dim1 plays a direct role in recognizing the 5′ splice site (Bertram et al. 2017). Dim2, an ortholog of DROL1, has also been identified in the human U12-dependent B complex (Bai et al. 2024). Although the imaging resolution was low and the mechanism underlying the interaction between Dim2 and the 5′ splice site remains unclear, Dim2 may recognize the 5′-terminal AU in a manner similar to Dim1.

Another gene, *CENATAC*, has been identified as a cause of human mosaic variegated aneuploidy (de Wolf et al. 2021). The CENATAC protein was originally reported to be a subunit of the U12-dependent spliceosome, as it coprecipitated with minor spliceosome-specific proteins and the U4atac and U6atac snRNAs. Comprehensive transcriptomic analysis of CENATAC-depleted cells revealed increased retention of U12-type introns, particularly those of the AT–AC subtype. Taken together, these findings indicate that CENATAC is required for the splicing of AT–AC introns, similar to DROL1, although it also participates in the splicing of some U12-type GT–AG introns.

CENATAC has also been identified in structural models of the U12-dependent spliceosomal B complex (Bai et al. 2024). In this complex, CENATAC binds to the 5′ cap of U4atac snRNA and to the loop region of U6atac snRNA. However, because CENATAC is not positioned adjacent to an intron, it remains unclear how this factor specifically acts on AT–AC introns.

The *Arabidopsis* ortholog of *CENATAC* has been named *TITAN LIKE* (*TTL*) because the mutant phenotype resembles that of the *titan* mutant, exhibiting abnormalities in embryo development (Lu et al. 2012). *TTL* encodes a protein containing a C2H2 domain, but the mechanism by which its loss or dysfunction leads to embryonic lethality remains unknown.

An interesting feature of the *CENATAC*/*TTL* gene family is that these genes contain a U12-type intron whose removal prevents the production of a functional protein (de Wolf et al. 2021). In vertebrates, *CENATAC* orthologs possess a U12-type intron with GT–AG termini that shares a 3′ splice site with a U2-type intron. Consequently, when splicing occurs at the U12-type 5′ splice site, the translational reading frame of the following exon is shifted, resulting in a nonfunctional protein (de Wolf et al. 2021). In plants, *TTL* orthologs typically contain AT–AC introns within coding exons. Moreover, splicing of these introns can lead to the loss of amino acid sequences, thereby also causing a frameshift (de Wolf et al. 2021; Lu et al. 2012). *TTL* undergoes alternative splicing to produce nine distinct transcripts, but only those retaining the AT–AC intron have been shown to complement the *ttl* mutant phenotype (Lu et al. 2012).

In this study, we investigated the function of *Arabidopsis* TTL in pre-mRNA splicing. To this end, we generated a viable TTL allele and identified a mutant, designated *ttl-142*, which carries a 42-nucleotide deletion in its first exon. The *ttl-142* mutant exhibited a mild morphological phenotype similar to that of *drol1* but showed a stronger suppression of AT–AC intron splicing. Moreover, unlike *drol1*, splicing of U12-type GT–AG introns was also reduced in *ttl-142*. Compared with wild-type (WT) *A. thaliana*, *ttl-142* displayed fewer differentially expressed genes (DEGs) than *drol1*, a finding consistent with its morphological phenotype. Taken together, these results indicate that TTL, like CENATAC, functions as a splicing factor required for the efficient processing of U12-type introns, particularly those of the AT–AC subtype. Furthermore, we speculate that the phenotypes of *drol1* and *ttl-142* plants may result partly from genetic defects due to AT–AC intron retention and partly from distinct splicing disruptions caused by each mutation.

## Results and Discussion

### ttl-142 is a weak allele of TITAN LIKE

Since *Arabidopsis TITAN LIKE* (TTL) is homologous to human *CENATAC*, it has been speculated to be required for the splicing of U12-type introns, particularly those of the AT–AC subtype. However, analysis of splicing patterns in the original *ttl* mutant has been difficult because of its embryonic lethality. To overcome this limitation and to generate viable *Arabidopsis* mutants for TTL, single guide RNAs (sgRNAs) were designed and introduced into WT (Col-0) plants. One transformant that received sgRNA1 was found to lack 42 nucleotides in the first exon of *TTL*; this mutant was designated *ttl-142* (Figures 1A and 1B). Plants homozygous for *ttl-142* were then phenotyped and found to be smaller than WT plants, displaying flattened cotyledons and dark green leaves (Figures 1C and 1D). Overall, these phenotypes were reminiscent of those of *drol1* mutants, although less severe (Figure 1E).

**Figure 1.**
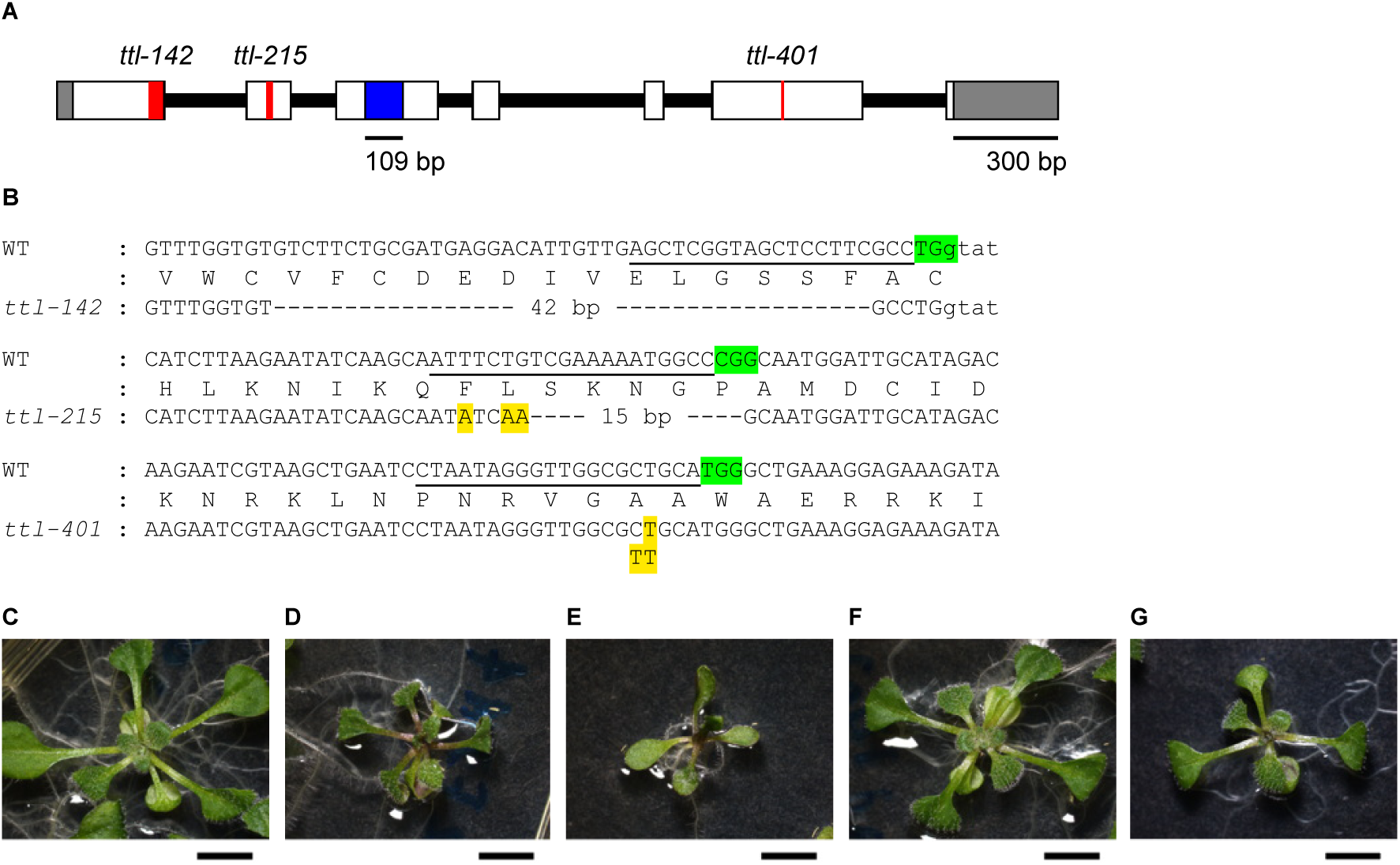
Generation of TTL mutants. (A) Structure of *TTL*. Exons are shown as boxes, with white regions indicating coding sequences and gray regions indicating UTRs. The blue box indicates the AT–AC intron located in exon 3, and the red boxes denote the *ttl-142, ttl-215*, and *ttl-401* mutations, respectively. (B) Nucleotide sequences surrounding the TTL genomic region targeted for genome editing. The WT nucleotide and deduced amino acid sequences are shown in the first and second lines, respectively. The sgRNA target sequence is underlined, and the protospacer adjacent motif is shown in green. Exonic and intronic regions are indicated in uppercase and lowercase letters, respectively. The sequences of the *ttl-142*, *ttl-215*, and *ttl-401* alleles generated by genome editing are shown below the WT sequence, and numbers within the sequences indicate the length of nucleotide deletions. (C–G) Phenotypes of WT, *ttl-142*, *drol1-1*, *TTLg/ttl-142*, and　*ttl-215*. All photographs were taken 13 days after germination. Scale bar = 5 mm.

To confirm that these phenotypes were caused by the *ttl-142* mutation, we introduced a DNA fragment containing the entire *TTL* gene, including 1 kb of the upstream region and 0.6 kb of the downstream region, into *ttl-142* plants. The resulting transformants exhibited a phenotype highly similar to that of the WT (Figure 1F), indicating that the phenotypes observed in *ttl-142* were indeed attributable to the *TTL* mutation rather than to off-target effects.

In addition to *ttl-142*, we obtained two other *TTL* mutants using different sgRNAs. A transformant that received sgRNA2 was found to lack 15 nucleotides in the second exon of *TTL*, and this mutant was designated *ttl-215* (Figures 1A and 1B). Homozygous *ttl-215* plants exhibited a phenotype that was slightly smaller than that of WT plants (Figures 1C and 1G). Another transformant that received sgRNA4 carried a thymidine insertion in the sixth exon and was designated *ttl-401* (Figures 1A and 1B). Among 104 offspring obtained by self-pollination of *ttl-401* heterozygotes, 29 were homozygous WT, 75 were heterozygous for *ttl-401*, and no *ttl-401* homozygotes were recovered. This segregation pattern is consistent with the embryo-lethal phenotype reported for the original *ttl* mutant (Lu et al. 2012).

Consistent with their shoot phenotypes, root growth was also impaired in the *ttl* mutants. Roots of *ttl-142* plants were 15–20% shorter than those of WT, whereas *ttl-215* roots were 20–45% shorter (Figure 2). By contrast, *drol1-1* seedlings germinated more slowly, and their roots were significantly shorter than those of WT when compared on the same day.

**Figure 2.**
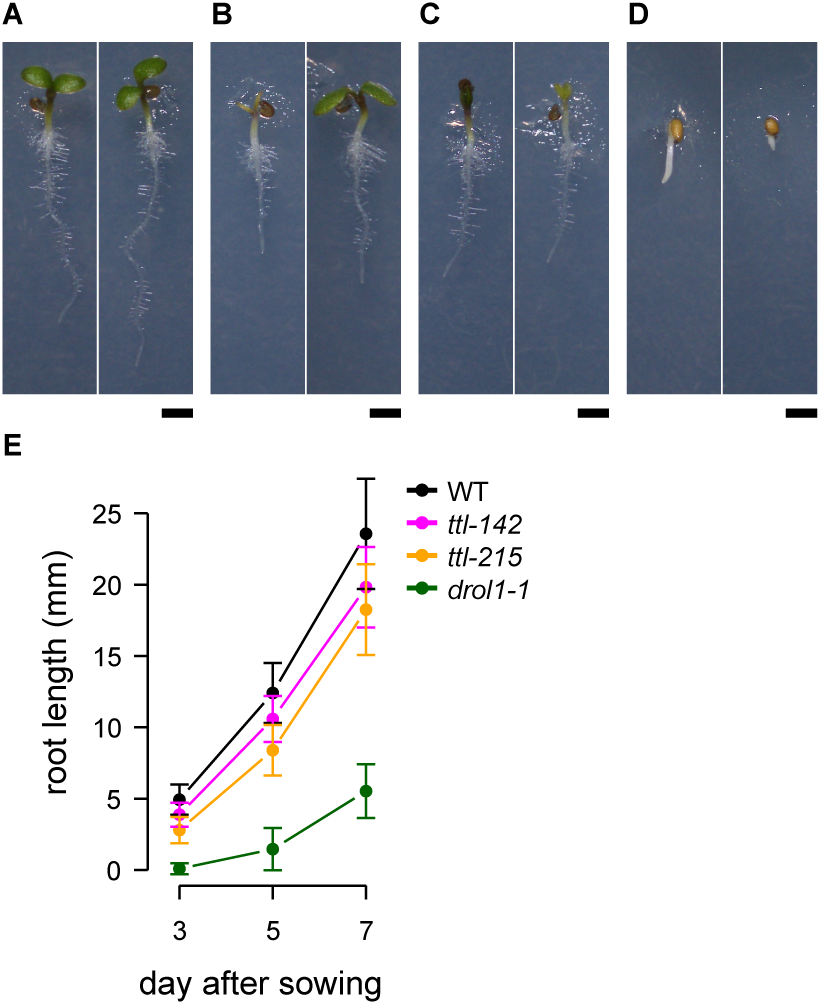
Root phenotypes and growth defects of *ttl* mutants. (A–D) Seedlings of 3-day-old WT, *ttl-142*, *ttl-215*, and *drol1-1*. Scale bar = 1 mm. (E) Root length of 3–7-day-old seedlings. Error bars represent standard deviation (n = 32–95).

Functional prediction indicated that the *ttl-142* mutation results in the production of a TTL protein lacking 14 amino acids. Notably, this deletion occurs within the second of the two C2H2-type zinc finger domains in TTL (Figure S1). In a previous structural study of the human minor spliceosome, the TTL homolog CENATAC was shown to use these two zinc fingers to bind the 5′ cap of U4atac and the central stem-loop of U6atac snRNAs (Figure S1; Bai et al. 2024). These findings suggest that the *ttl-142* mutation disrupts the binding of TTL to the U4atac/U6atac di-snRNP complex. The *ttl-215* mutation is located in a less conserved region following the second C2H2 domain. Given that *ttl-215* exhibits phenotypes similar to those of *ttl-142*, this mutation is also likely to impair TTL function in a similar manner.

In addition to the C2H2 motif, the *CENATAC***/***TTL* gene family contains several other conserved motifs. Four motifs of unknown function (M1–M4) are conserved in the C-termini of metazoan homologs (de Wolf et al. 2021), but *TTL* lacks a region with high similarity to M2 (Figure S1D). In human patients, a mutation altering a splice site results in the loss of M3 and M4 (de Wolf et al. 2021), and the *ttl-401* mutation in *Arabidopsis* similarly disrupts M3 and eliminates M4 due to a frameshift. These results indicate that these motifs are essential for CENATAC/TTL function. In the three-dimensional structure predicted by AlphaFold3 (Abramson et al. 2024), these motifs form part of an α-helix (Figures S1B and S1C); however, they have not yet been experimentally confirmed. The region between the C2H2 domain and M1 is an intrinsically disordered domain, predicted to have no defined structure. Future studies should focus on elucidating the role of these motifs in *TTL* function and their contribution to splicing activity.

### RNA-Seq analysis of splicing in ttl-142

RNA extracted from whole 3-day-old *ttl-142* seedlings was subjected to RNA-Seq analysis. The resulting transcriptomic dataset was compared with those from WT plants and the *drol1-1* mutant (Suzuki et al. 2022). Splicing patterns were analyzed using ASpli (Estefania et al. 2021), and the percentage of intron retention (PIR) was calculated. As shown in Figure 3A, most PIR values were similar between WT and *ttl-142* plants; however, 479 introns exhibited significant differences (false discovery rate [FDR] < 0.01; red and blue dots in Figure 3A). Among these, 60 introns (13%; red dots in Figure 3A) began with AT and ended with AC—that is, putative AT–AC introns. This proportion was markedly higher than that in the *Arabidopsis* genome as a whole (0.06%; *P* < 2.2 × 10^−16^, chi-square test). Furthermore, among the 479 introns identified, PIRs of approximately half of the non–AT–AC introns were significantly increased, whereas the other half were decreased in *ttl-142*. Conversely, the PIRs of nearly all AT–AC introns increased, indicating that splicing of AT–AC introns was specifically suppressed in the *ttl-142* mutant. Genes containing AT–AC introns whose PIRs were altered in *ttl-142* are listed in Table 1.

**Figure 3.**
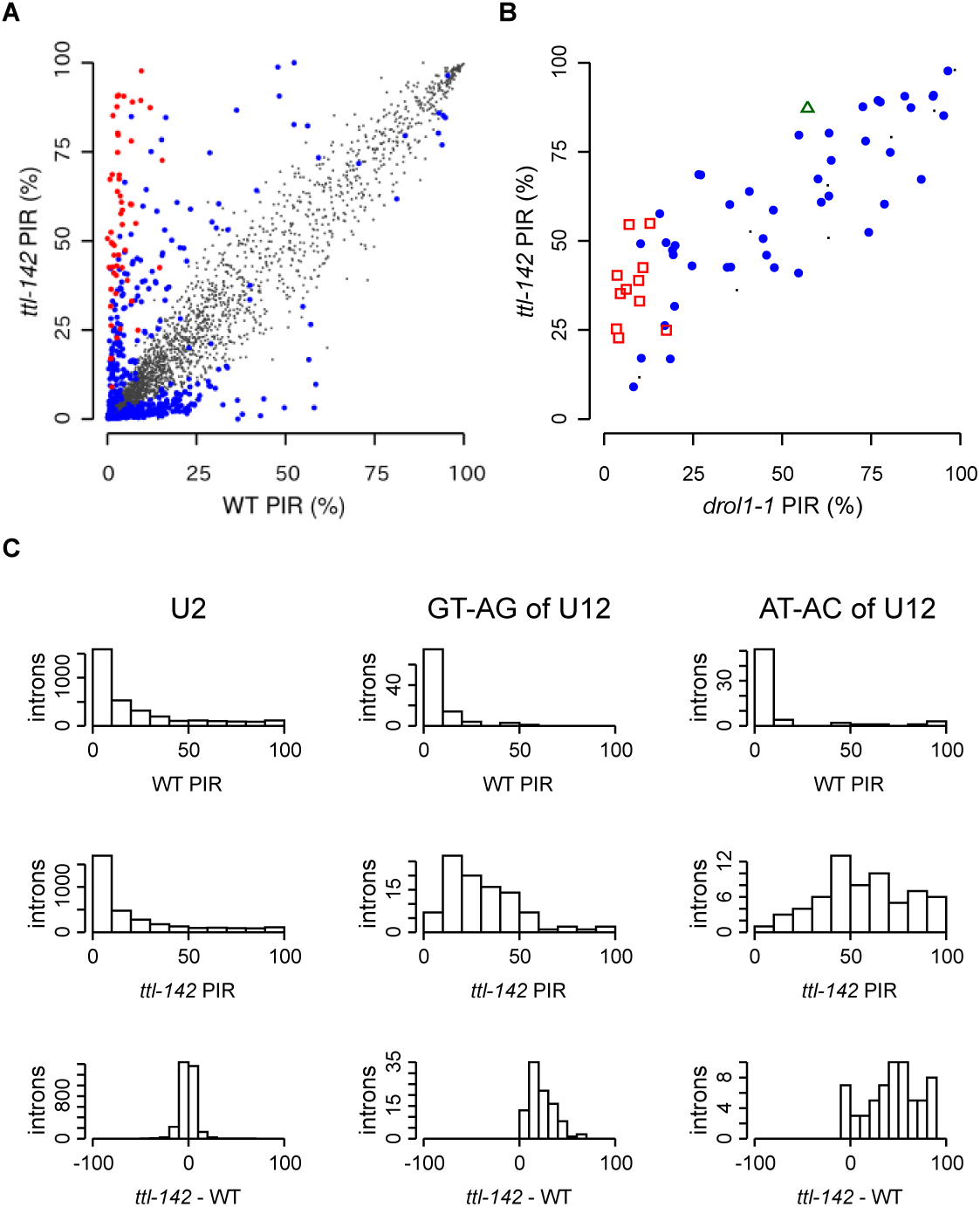
Global intron retention in the *ttl-142* mutant. (A) Scatter plot of the PIR in WT and *ttl-142* plants. A total of 3,387 introns were analyzed using ASpli (Estefania et al. 2021). AT–AC introns and other intron types (mainly GT–AG) with FDR < 0.01 are plotted as large red and blue dots, respectively; small dots represent introns without significant changes. (B) Comparison of PIR between *ttl-142* and *drol1-1*. PIR values for all AT–AC introns in *ttl-142* and *drol1-1* are plotted on the vertical and horizontal axes, respectively. Red squares, green triangles, and large blue dots represent AT–AC introns with FDR < 0.01 (relative to WT) in *ttl-142*, *drol1-1*, and both mutants, respectively. Small dots denote AT–AC introns with no significant change in PIR. (C) Histograms of PIR distributions. From left to right, columns show U2 introns, U12-type GT–AG introns, and U12-type AT–AC introns grouped by PIR values. The first and second rows display observed PIR values in WT and *ttl-142*, respectively, and the third row shows the differences in PIR (*ttl-142* minus WT). U12-type introns were downloaded from ERISdb (Szcześniak et al. 2013).

**Table 1.**
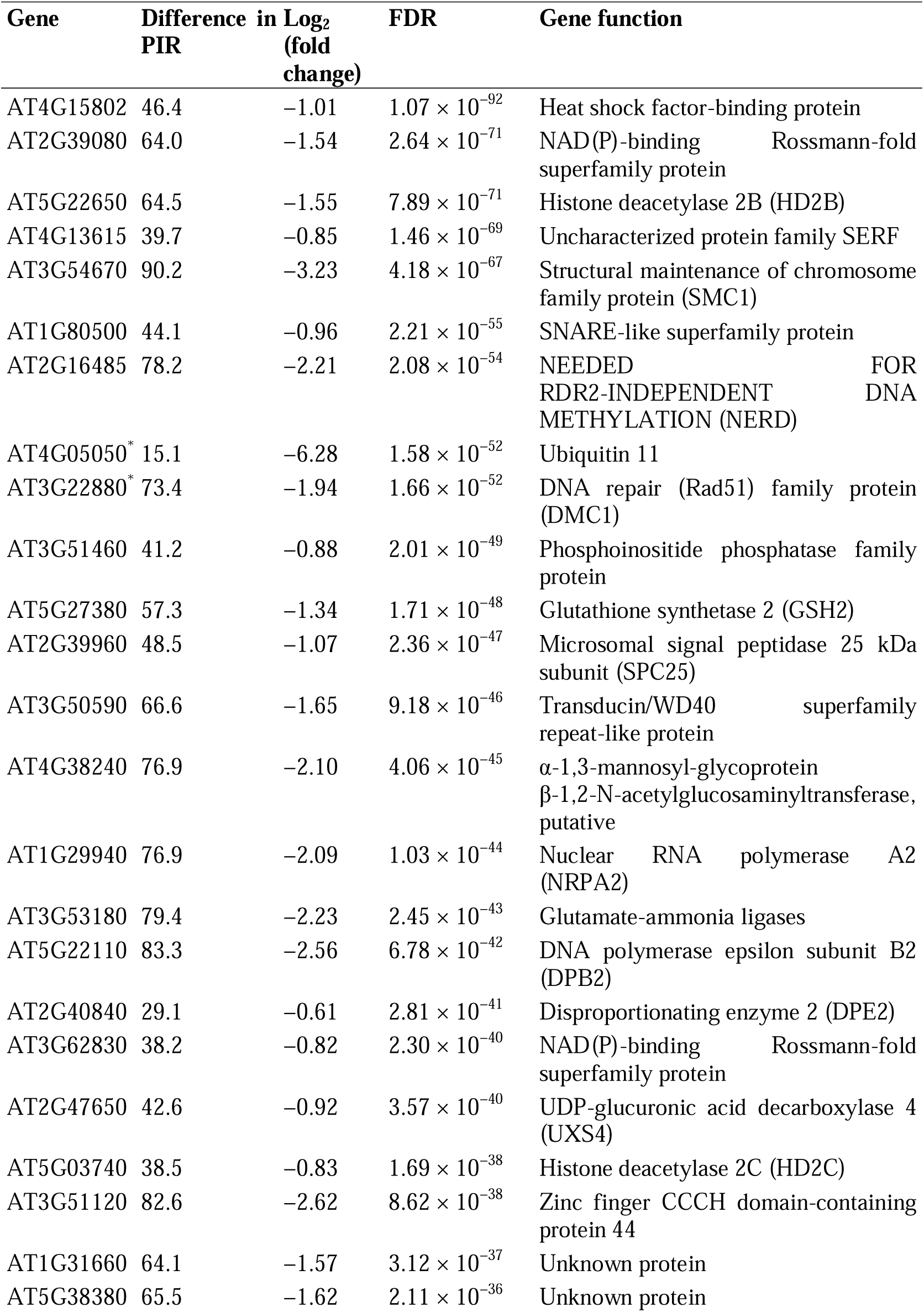
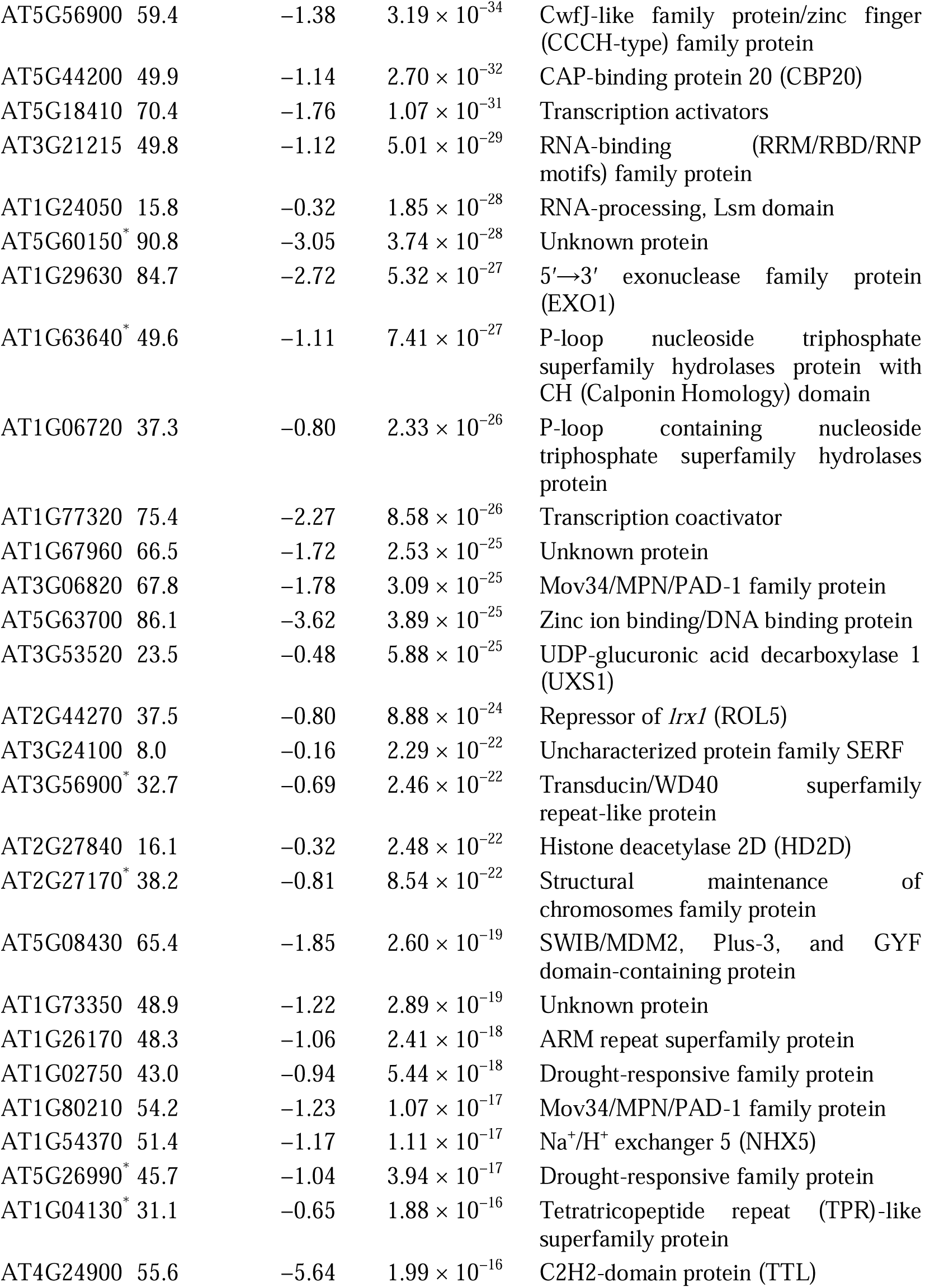
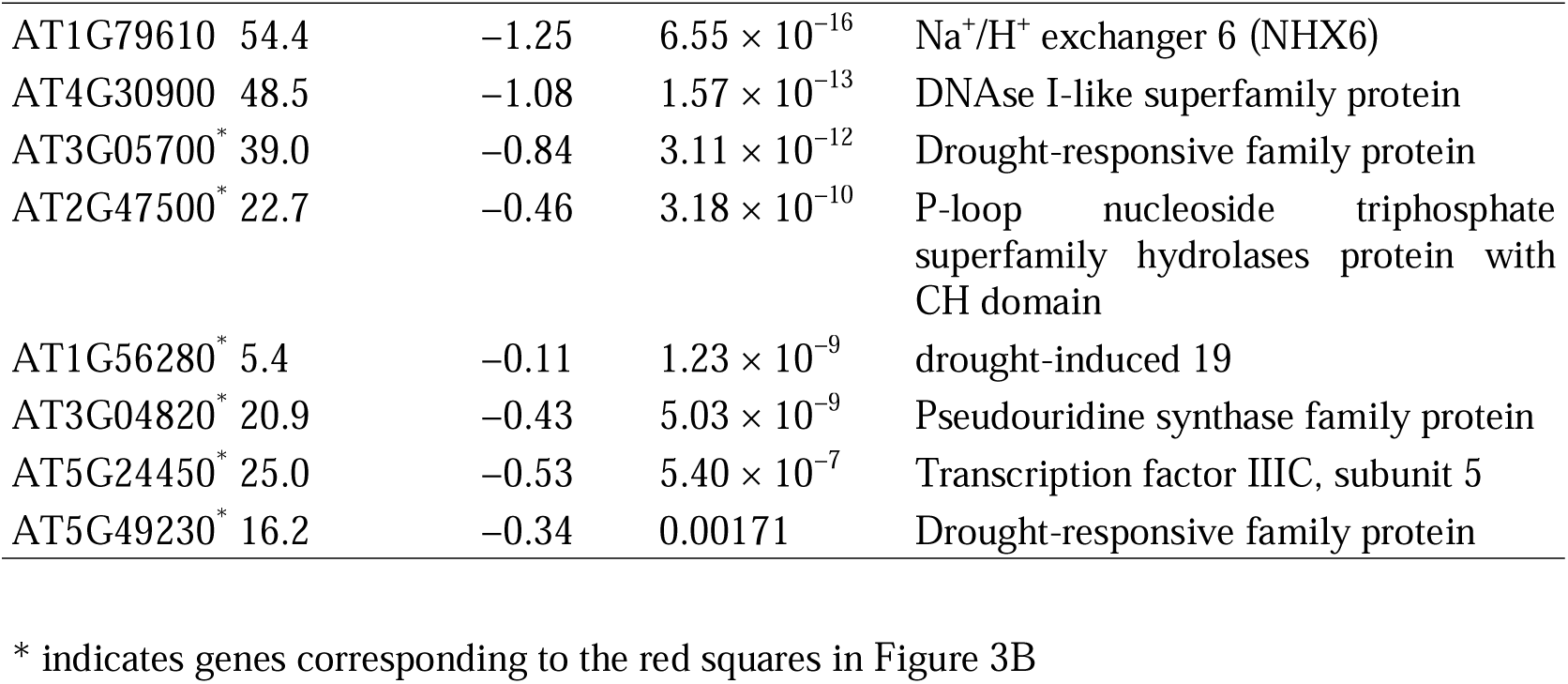
Genes with AT–AC-type introns showing significant intron retention in *ttl-142* mutants.

The splicing patterns of AT–AC introns were next compared between the *ttl-142* and *drol1-1* mutants (Figure 3B). Among the 79 AT–AC introns analyzed, 47 showed significant changes in PIR in both *ttl-142* and *drol1-1* relative to the WT (FDR < 0.01; blue dots in Figure 3B). On average, PIR increased by 9% in *ttl-142* (*P* < 0.001, Student’s *t*-test). Eighteen AT–AC introns exhibited significantly increased PIR values only in *ttl-142* compared with WT (red squares in Figure 3B), whereas only one intron showed a significant increase in *drol1-1* alone (green triangle in Figure 3B). Taken together, these results indicate that splicing of AT–AC introns was more strongly suppressed in *ttl-142* than in *drol1-1*. This finding is inconsistent with the observation that *ttl-142* exhibits a milder morphological phenotype than *drol1-1* and suggests that the severe phenotype of *drol1-1* is not solely attributable to inhibition of AT–AC intron splicing.

In general, AT–AC introns are spliced by the U12-dependent spliceosome, which also processes introns with GT–AG termini. We next examined the dependency of each intron type on TTL function. The U2-dependent spliceosome, the major spliceosome in eukaryotes, is responsible for splicing most introns. As shown in Figure 3C (left column), the PIR values of introns spliced by the U2-dependent spliceosome were zero or nearly zero in both WT and *ttl-142* plants. Moreover, no significant differences in PIR were observed between WT and *ttl-142* for U2-type introns (Figure 3C, lower left), indicating that TTL is not required for U2-dependent splicing. Conversely, the PIR values of most U12-type GT–AG introns were near zero in WT plants but increased in *ttl-142* (Figure 3C, middle column). On average, the difference in PIR between *ttl-142* and WT for these introns was 24%, which was significantly different from zero (*P* < 2.2 × 10^−16^, Student’s *t*-test). Furthermore, the PIR values of U12-type AT–AC introns were significantly higher in *ttl-142* than in WT plants (*P* < 2.2 × 10^−16^, Student’s *t*-test), with an average difference of 45% (Figure 3C, right column). In *ttl-142* plants, PIR values of U12-type AT–AC introns were significantly greater than those of U12-type GT–AG introns (*P* < 1.3 × 10^−8^, Student’s *t*-test). Taken together, these results indicate that TTL is required for U12-dependent intron splicing, particularly for AT–AC introns. This finding is consistent with previous results for human CENATAC but differs from DROL1, which is required exclusively for the splicing of AT–AC introns.

Structural analysis of the human minor spliceosome has revealed that CENATAC binds to the U4atac/U6atac di-snRNP (Bai et al. 2024). A previous biochemical study also shown that CENATAC primarily associates with the U4atac/U6atac di-snRNP (de Wolf et al. 2021). These findings are consistent with the observation that when CENATAC or TTL function is impaired, splicing of all U12-type introns is suppressed. However, because CENATAC is positioned far from the 5′ end of the intron within the spliceosome, this localization cannot explain the observed differences in dependency between U12-type GT–AG and AT–AC introns. Conversely, Dim1, a homolog of DROL1, is known to be located adjacent to the 5′ end of the intron (Bertram et al. 2017; Zhan et al. 2018). Functional analysis of DROL1 has shown that it regulates the specific binding of U5 snRNPs to the 5′ end of the intron (Suzuki et al. 2025). Further analysis will be required to elucidate why U12-type AT–AC introns are more dependent on TTL.

### RT-PCR analysis of AT–AC intron retention in ttl-142

Figure 4A shows the compiled RNA-Seq reads derived from WT and *ttl-142* seedlings mapped to the *TTL* gene. Short reads from WT seedlings primarily mapped to exons, with minimal coverage in introns other than the fourth intron (Figure 4A, upper panel). Overall, *ttl-142* exhibited a pattern similar to WT except for the 42-nucleotide deletion in the first exon (red arrowhead) and the AT–AC intron located in the third exon. The PIR for this intron was 44% in WT but reached 100% in *ttl-142*. This result was further validated via reverse transcription PCR (RT-PCR). RT-PCR was performed using primers designed upstream of the AT–AC intron and on exon 4 (Figure 4B). In WT plants, three peaks were detected, one of which, located around 250 bp, could not be identified. The remaining two peaks were assigned to mRNAs with or without the AT–AC intron (Figure 4C). In contrast, in the *ttl-142* mutant, the peak representing the AT–AC intron-containing mRNA was predominant, whereas the peak corresponding to the spliced product was barely detectable (Figure 4D).

**Figure 4.**
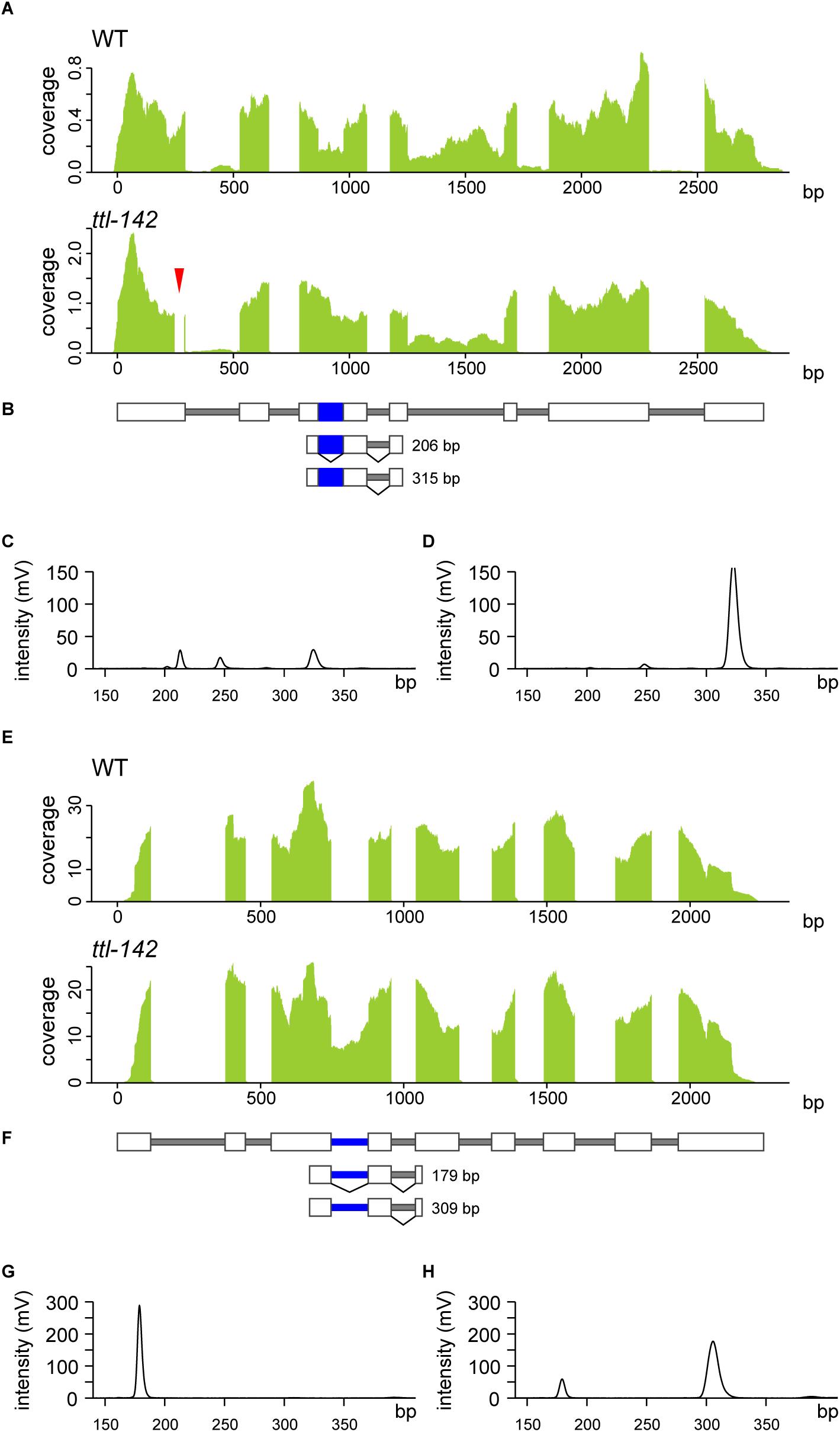
RNA-Seq and RT-PCR analyses of AT–AC intron retention in *TTL* and *HD2B* in WT and *ttl-142*. (A) RNA-Seq analysis of *TTL*. Total RNA was extracted from 3-day-old WT (upper) and *ttl-142* (lower) seedlings, followed by cDNA library construction and next-generation sequencing. All reads obtained were aligned to the *TTL* gene sequence. The x-axis represents reference sequences, and the y-axis indicates short-read coverage. The red arrowhead marks the 42-nucleotide deletion in *ttl-142*. (B) Structure of the *TTL* gene. Gray and white boxes represent introns and exons, respectively, and blue boxes indicate AT–AC-type introns. Expected RT-PCR products shown in (C) and (D) are also indicated, along with their expected fragment sizes. (C, D) Microchip electrophoresis of DNA fragments amplified via RT-PCR from (C) WT and (D) *ttl-142* RNA. The x-axis represents DNA fragment size (bp), and the y-axis indicates fluorescence intensity corresponding to DNA abundance. (E-H) Corresponding analyses for *HD2B*.

The *TTL* gene contains an AT–AC intron within its third exon that encodes part of the functional protein. In WT plants, approximately half of TTL transcripts retain the AT–AC intron and encode functional proteins, whereas the remainder are spliced to remove this intron. This balance may be maintained through a negative feedback mechanism, as TTL itself promotes splicing of the AT–AC intron. In our RNA-Seq dataset, the abundance of TTL transcripts in *ttl-142* was approximately 2.4-fold higher than that in WT, supporting the possibility that TTL expression is regulated by negative feedback.

The histone deacetylase 2B (*HD2B*) gene belongs to the *HD2* gene family, which encodes plant-specific histone deacetylases (Hollender and Liu 2008). Notably, its third intron is an AT–AC intron. In WT seedlings, all introns were fully spliced, whereas in *ttl-142*, 65% of the AT–AC introns were retained (Figure 4E). Additionally, DNA fragments containing AT–AC introns were undetectable in WT transcriptomes but were markedly more abundant in *ttl-142*, as confirmed via RT-PCR (Figures 4F–4H). Finally, no significant difference was observed in *HD2B* expression levels between WT and *ttl-142*.

A similar analysis was performed on five genes (*HSI2*, *AT1G79880*, *AT5G06620*, *AT4G23330*, and *AT1G18090*) that contain U12-type GT–AG introns. In each case, U12-type intron–specific retention was increased in *ttl-142* compared with WT and *drol1-1*, confirming that *TTL* is required for U12-dependent splicing (Figures S2–S6).

### DEGs in the ttl-142 mutant

We next compared genome-wide gene expression patterns between WT and *ttl-142* plants. DEGs were identified using ASpli (Estefania et al. 2021) with an FDR threshold of <1 × 10^−4^. Compared with WT, 295 genes were significantly upregulated and 657 were downregulated in *ttl-142* (Figure 5A). In contrast, 1402 genes were upregulated and 1376 were downregulated in *drol1-1* compared with WT (Figure 5B). Overall, *drol1-1* exhibited a larger number of DEGs than *ttl-142*, which may account for its stronger developmental phenotype; however, this difference could not be explained by the degree of intron retention.

**Figure 5.**
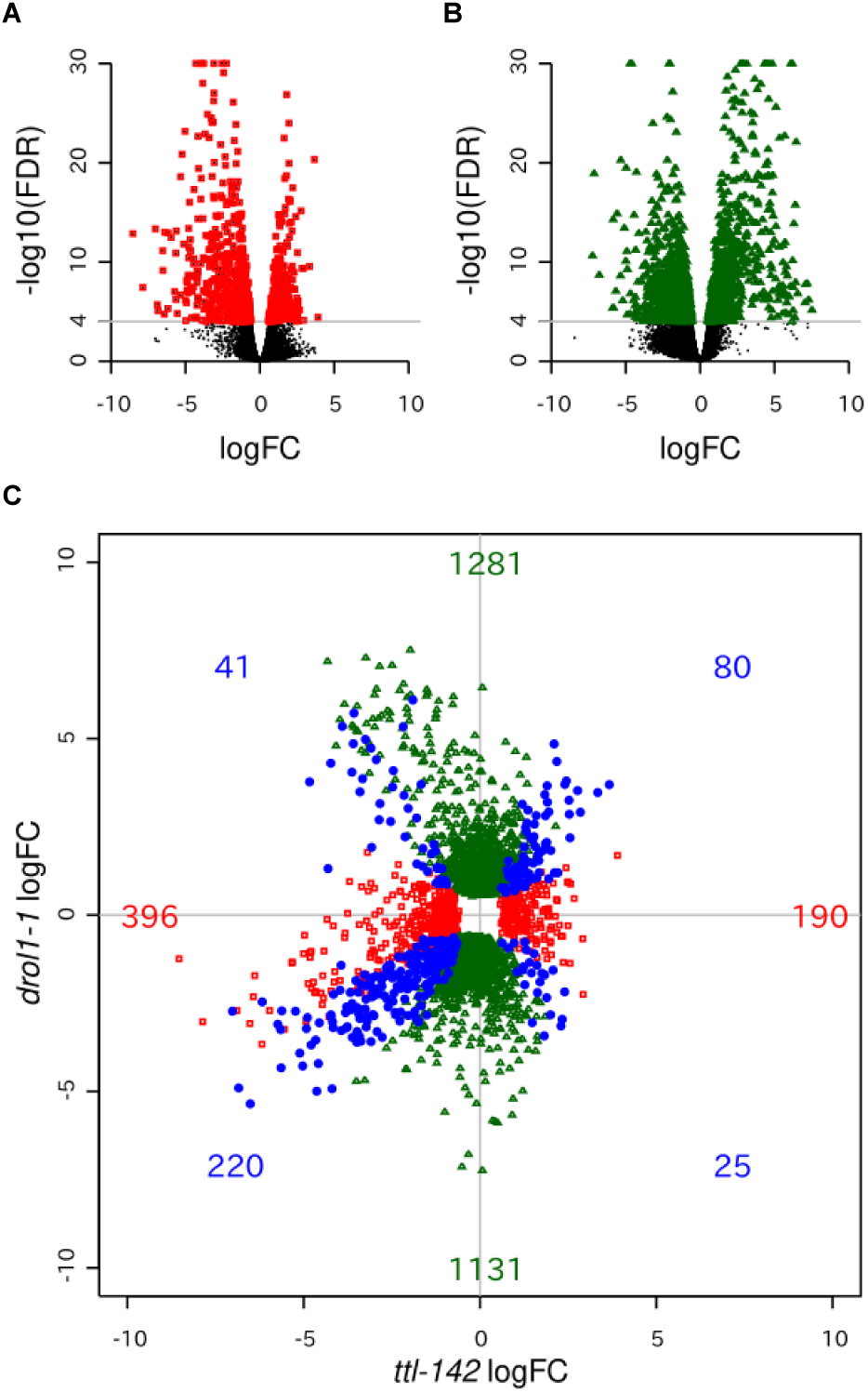
Comparison of DEGs in *ttl-142* and *drol1-1*. (A) Volcano plot showing DEGs in *ttl-142* relative to WT. Genes with FDR < 1 × 10^−4^ are highlighted with red squares. FDR values smaller than 1 × 10^−30^ were rounded to 1 × 10^−30^ for plotting. (B) Volcano plot showing DEGs in *drol1-1* relative to WT. Genes with FDR < 1 × 10^−4^ are highlighted with green triangles. (C) Log_2_ fold changes in gene expression levels in *ttl-142* and *drol1-1* were plotted on the vertical and horizontal axes, respectively. Red squares, green triangles, and blue dots represent genes with FDR < 1 × 10^−4^ in *ttl-142*, *drol1-1*, and both mutants, respectively. Numbers within the plot indicate the number of genes in each group.

When comparing the *ttl-142* and *drol1-1* DEG datasets, we identified 80 genes that were upregulated and 220 that were downregulated in both mutants (Figure 5C, blue circles). However, 66 genes exhibited opposing expression changes between *ttl-142* and *drol1-1* (Figure 5C, blue circles), indicating that the overall gene expression patterns of *ttl-142* and *drol1-1* are distinct.

Previous studies reported that gene ontology (GO) terms such as “lipid storage,” “response to abscisic acid,” “response to water deprivation,” and “response to salt stress” were over-represented among genes upregulated in *drol1* relative to WT, whereas “mitotic cell cycle transition” and “regulation of cyclin-dependent protein serine/threonine kinase activity” were over-represented among genes downregulated in *drol1* (Suzuki et al. 2022). In this study, a similar GO analysis of *ttl-142* revealed no significantly enriched ontologies among genes whose expression levels were either increased or decreased. Moreover, no significantly enriched GO terms were identified among genes that were commonly up- or downregulated in both *drol1* and *ttl-142*. Taken together, these findings indicate that the upregulation of abscisic acid (ABA)-related genes and the downregulation of cell division–related genes in *drol1* are specific to that mutant and are not necessarily associated with inhibition of AT–AC intron splicing.

Although *drol1* and *ttl-142* display similar phenotypes in cotyledon shape and leaf color, the *drol1* phenotype is more severe (Figure 1). Previously, the *drol1* phenotype was attributed to splicing suppression specific to AT–AC introns (Suzuki et al. 2022). However, in *ttl-142*, where splicing suppression is even more pronounced than in *drol1*, the phenotype is milder. This indicates that the *drol1* phenotype cannot be explained by splicing suppression alone. Additionally, *drol1* exhibits high sensitivity to ABA, with significant upregulation of ABA pathway–related genes relative to WT (Suzuki et al. 2022). Taken together, these findings suggest that the loss of DROL1 specifically impairs a particular stage of splicing, which in turn activates the ABA response.

### Subcellular localization of the TTL protein

We next examined the subcellular localization of the TTL protein. A DNA fragment containing 1 kbp upstream and the full coding sequence of TTL was amplified from the *Arabidopsis* genome and fused at its C-terminus to GFP. Introduction of this recombinant construct (TTLn-GFP) into *ttl-142* rescued the mutant phenotype (Figure 6A), but GFP fluorescence was not detected (10 independent T2 lines were examined). RNA was extracted from the transformants and reverse-transcribed using a GFP-specific primer. RT-PCR revealed that approximately half of the AT–AC intron was spliced (Figure 6B). We speculated that alternative splicing of the AT–AC intron reduces the abundance of transcripts encoding a functional TTL protein.

**Figure 6.**
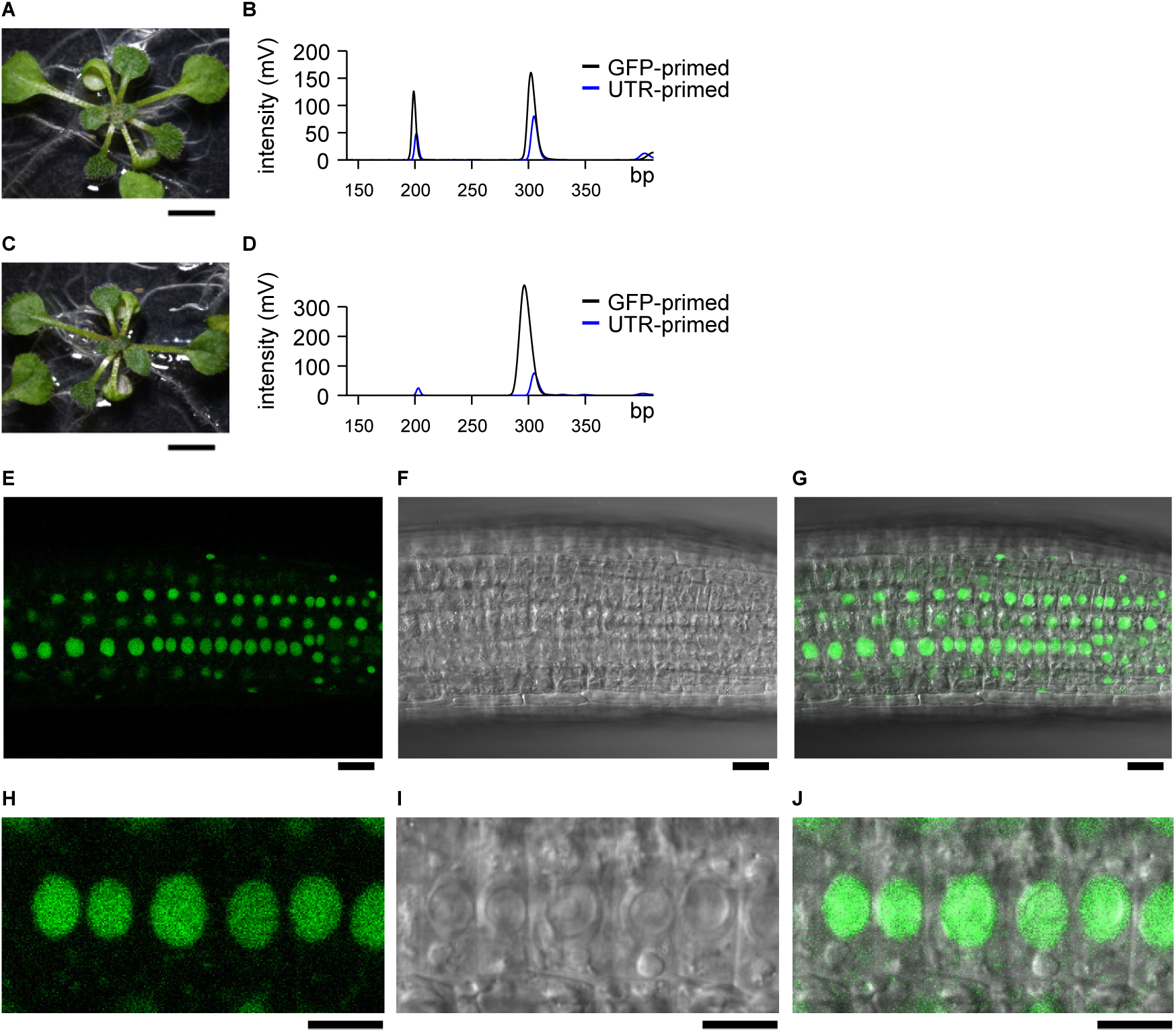
Subcellular localization of TTL in root cells. (A and C) Phenotypes of *TTLn-GFP* / *ttl-142* and *TTLnmI-GFP* / *ttl-142*. All photographs were taken 13 days after germination. Scale bar = 5 mm. (B and D) RT-PCR analysis of TTL splicing. RNA extracted from (B) *TTLn-GFP* / *ttl-142* and (D) *TTLnmI-GFP* / *ttl-142* was reverse-transcribed using primers for GFP (black line) or for the 3′ UTR of TTL (blue line). Subsequent PCR and electrophoresis were performed as described in Figure 4C. (E-G) Root of a WT transformant expressing TTLnmI–GFP. Shown are (E) fluorescence from TTLnmI–GFP, (F) a differential interference contrast image, and (G) the merged image. Scale bar = 20 µm. (H–J) Magnified view of (E–G). Scale bar = 20 µm. All images were acquired using an LSM 710 AxioObserver confocal microscope (Zeiss, Jena, Germany).

To test this, we introduced 23 base substitutions into the AT–AC intron without altering the amino acid sequence (Figure S7). The modified gene was designated *TTLnmI*. When GFP was fused to *TTLnmI* and expressed in *ttl-142*, the phenotype was restored (Figure 6C). RT-PCR confirmed that the modified AT–AC intron was no longer spliced (Figure 6D). Strong GFP fluorescence was observed in root (Figures 6E–6J), and the signal was localized to the nucleus, consistent with previous results obtained using protoplasts (Lu et al. 2012).

The fluorescent intensities of TTLn-GFP and TTLnmI-GFP were not correlated with the amounts of their transcripts, suggesting that TTLn-GFP mRNAs retaining the AT–AC intron are not used for translation. These results are consistent with our previous report showing that the retained AT–AC intron inhibits mRNA export to the cytoplasm (Suzuki et al. 2025).

Because *TTL* mRNA levels were increased in *ttl-142* mutants, we propose that TTL expression is regulated through a negative feedback loop (de Wolf et al. 2021). Furthermore, GFP fused to TTL was not expressed in transformants, whereas GFP fluorescence was detected when mutations were introduced into the AT–AC intron. Given that the AT–AC intron within the TTL exon encodes part of the functional protein and that TTL itself promotes splicing of this intron, we suggest that TTL protein abundance is controlled by a negative feedback mechanism mediated by AT–AC intron splicing. Such splicing-dependent autoregulation may represent a conserved feature of the *CENATAC***/***TTL* gene family.

## Conclusion

Arabidopsis *TTL* was originally isolated as a recessive embryonic lethal gene (Lu *et al*., 2012), which substantially hindered analysis of its role in pre-mRNA splicing. Using CRISPR-Cas9, we therefore generated two weak *TTL* alleles, *ttl-142* and *ttl-215*. RNA-Seq analysis showed that splicing of U12-type introns was suppressed in *ttl-142*, with AT–AC introns being particularly affected. This phenotype resembles that observed in human patients and in CENATAC-depleted cells (de Wolf *et al*., 2021), indicating that the CENATAC/*TTL* gene family constitutes a conserved factor required for efficient splicing of U12-type introns, especially those of the AT–AC subtype.

Although both *ttl-142* and *drol1* exhibit defects in AT–AC intron splicing and share similar developmental abnormalities, the severity of their phenotypes correlates with changes in gene expression rather than with the extent of splicing impairment. This suggests that defective splicing of AT–AC introns does not directly determine specific phenotypic outcomes. Instead, the distinct phenotypes of *ttl-142* and *drol1* are likely caused by the loss of their respective protein functions. We have ioslated *drol1* suppressors and demonstrated that their developmental phenotypes were indistinguishable from those of the WT, although the defects in AT-AC intron splicing were not fully restored (Suzuki et al. 2025). This result further indicates that the phenotypes of *ttl-142* and *drol1* are not caused solely by splicing impairment.

Expression of the *TTL* was increased in the *ttl-142* mutant, suggesting that *TTL* abundance may be regulated through a negative feedback loop (de Wolf *et al*., 2021). Consistent with this idea, TTL–GFP was not detected in transformants expressing the wild-type construct, whereas strong GFP fluorescence was observed when mutations were introduced into the AT–AC intron. Because the AT–AC intron within the *TTL* exon encodes part of the functional protein and *TTL* itself promotes splicing of this intron, these results indicate that *TTL* protein levels are regulated by a negative feedback mechanism mediated by AT–AC intron splicing. Such splicing-dependent autoregulation appears to represent a common feature of the CENATAC/*TTL* gene family.

The CENATAC/*TTL* family contains two C2H2 domains at the N-terminus and a conserved sequence of unknown function at the C-terminus, connected by an intrinsically disordered region. CENATAC has been shown to bind U4atac and U6atac snRNAs via its C2H2 domain (Bai et al. 2024). The *ttl-142* allele carries a deletion within one C2H2 domain, which likely reduces its ability to bind the 5′ cap of U4atac snRNA, yet *ttl-142* remains viable. This observation suggests that *TTL* binding to U4atac snRNA may be partially retained or dispensable for viability. In contrast, the mutation identified in humans eliminates the conserved C-terminal motif, and the recessive lethal allele *ttl-401* similarly lacks this region. Future studies will be required to determine how these structural features contribute to U12-dependent splicing and to fully elucidate the molecular function of *TTL*.

## Materials and Methods

### Plant material and growth conditions

*A. thaliana* (L.) Heynh. ecotype Columbia (Col-0) was used as the WT. The *drol1-1* mutant was characterized previously (Suzuki et al. 2018). Unless otherwise indicated, seeds were surface-sterilized and germinated on 0.3% (w/v) gellan gum plates containing half-strength Murashige and Skoog medium (Wako, Japan) supplemented with 2% (w/v) sucrose, 0.05% (w/v) 2-(N-morpholino)ethanesulfonic acid (pH adjusted to 5.7 with KOH), 100 mg/L myo-inositol, 10 mg/L thiamine hydrochloride, 1 mg/L nicotinic acid, and 1 mg/L pyridoxine hydrochloride. Plant growth conditions were as described previously (Suzuki et al. 2018).　For root observation and RNA extraction, seeds were surface-sterilized and plated on 1.5% (w/v) agar medium supplemented with 1% (w/v) sucrose. Plates were placed vertically in a growth chamber.

### Gene cloning and plasmid construction

Gene-editing constructs were prepared using the pKI1.1R vector and combinations of specific oligonucleotides (Tsutsui and Higashiyama 2016).

DNA fragments for TTLg were amplified from *Arabidopsis* genomic DNA using the attB1_TTLp and attB2_TTLt primers. The amplified fragments were ligated to a pDONR vector that had been inverse-PCR-amplified with the attL1 and attL2 primers, using an In-Fusion Cloning Kit (Takara, Tokyo, Japan). The resulting plasmid was designated pDONR-TTLg. The TTLg insert was subsequently transferred into pFAST-G01 (Shimada et al. 2010) via LR recombination using Gateway cloning (Thermo Fisher Scientific, MA, USA), and the resulting construct was used for complementation of *ttl-142*.

For the expression of GFP-fused TTL, pDONR-TTLg was used as a template for inverse PCR with the attB2_TTL and attL2 primers, and the resulting PCR product was circularized using an In-Fusion Cloning Kit to generate pDONR-TTLn. This plasmid was then used as a template for inverse PCR with TTL_mIS and TTL_mIA primers, and the self-ligated product was designated pDONR-TTLnmI. Finally, inserts were transferred into the pGWB550 binary vector (Nakagawa et al. 2007), which was subsequently used for transformation. All oligonucleotides used in this study are listed in Table S1.

### RNA extraction

Seedlings were harvested at the time points specified in the Results section, flash-frozen in liquid nitrogen, and ground into a fine powder using a pestle and mortar. Total RNA was extracted from the powdered tissue using the NucleoSpin RNA Plant Kit (Takara, Shiga, Japan), following the manufacturer’s instructions.

### RNA-Seq analysis

Library preparation, sequencing, and read trimming were performed using the same protocols as described in our previous study (Suzuki et al. 2022). The obtained reads were aligned to the genomic sequence using HISAT2 (Kim, Paggi, et al. 2019) with default parameters. PIR values for introns, as well as log(fold change) and FDR values for both introns and genes, were calculated using ASpli (Estefania et al. 2021).

### RT-PCR and microchip electrophoresis

For RT-PCR, cDNA was synthesized from 400 ng of total RNA using 100 U of SuperScript III reverse transcriptase (Thermo Fisher Scientific, MA, USA) and 1 pmol of each of antisense primers in a total reaction volume of 10 µL. The synthesized cDNA was then amplified via PCR using MightyAmp (Takara, Shiga, Japan) for 30 cycles. PCR products were diluted fivefold with water and analyzed using the MultiNA microchip electrophoresis system (Shimadzu, Kyoto, Japan). All primers used are listed in Table S1.

### Microscopy

The roots of seedlings were grown vertically for 4 days on medium containing 1% sucrose and 1.5% agar. Roots were then observed using an LSM 710 AxioObserver confocal microscope (Zeiss, Jena, Germany).

### Statistical analyses

All statistical analyses were performed using R version 4.3.3 (https://www.r-project.org/). Specifically, the t.test and chisq.test functions were used to conduct Student’s *t*-tests and chi-square tests, respectively.

## Data availability

The data underlying this article are available in the DNA Data Bank of Japan under accession number PRJDB20300. Data for WT and *drol1* were obtained under accession number PRJDB9966. Arabidopsis *ttl* mutants will be made available following publication through the RIKEN BioResource Research Center (https://epd.brc.riken.jp/en/). Plasmids generated in this study are available from the corresponding author upon reasonable request.

## Acknowledgments

We thank Ayami Furuta and Mine Morimoto for valuable technical assistance. We also thank Enago (www.enago.jp) for English language review.

## Author contributions

T.S. conceived the study; T.S. and T.N. designed the experiments; T.S., T.N., J.M., N.I., H.I., and T.K. performed the experiments; T.S. analyzed the data; and T.S. wrote the manuscript.

