## Supplementary Tables for "*Arabidopsis TITAN LIKE* is required for U12-type intron splicing, especially of AT–AC subtypes"

**Table S1**

**Oligo DNA for sgRNAs**

| | |
|---|---|
| TTL_2271S+ATTG | ATTGAGCTCGGTAGCTCCTTCGCC |
| TTL_2290A+AAAC | AAACGGCGAAGGAGCTACCGAGCT |
| TTL_2581S+ATTG | ATTGATTTCTGTCGAAAAATGGCC |
| TTL_2600A+AAAC | AAACGGCCATTTTTCGACAGAAAT |
| TTL_4043S+ATTG | ATTGCTAATAGGGTTGGCGCTGCA |
| TTL_4062A+AAAC | AAACTGCAGCGCCAACCCTATTAG |

**Primers for cloning**

| | |
|---|---|
| attB1_TTLp | ACAAAAAAGCAGGCTGGCCAACAGTTACCCGAAC |
| attB2_TTLt | ACAAGAAAGCTGGGTTTTGCTCACCAGAAACAGTC |
| attB1_TTL | ACAAAAAAGCAGGCTATGAAGAAACCATCGAAAAA |
| attB2_TTL | ACAAGAAAGCTGGGTACTCACCACTCTCTCTTCG |
| attL1 | AGCCTGCTTTTTTGTACAAAGTTGG |
| attL2 | ACCCAGCTTTCTTGTACAAAGTTGG |
| TTL_mIA | CCTTCTTGATCCGGTCCATCGTCTCGAACGCCAGCTTCGTG TGAATGTCATTAGATGTCCCAGATAGCTG |
| TTL_mIS | ACCGGATCAAGAAGGTGCCGGCCCACCATATAAATTCGTAT AAGTCGAATGATGTGATGCCGTTGCAGTA |

**Primers for RT-PCR analysis**

| | |
|---|---|
| TTL_2818S | ATGCATCTTTTGAAGGGTCCTG |
| TTL_3229A | GTTGCCCGAGTAGTCCTTGCT |
| HD2B_3114A | GACAACAGCTGCTCCAGCATT |
| HD2B_2721S | AAGAGTTTGAGCTTTCACACAGCG |
| HSI2_U12_S | CACTTGCTTCCGCGGTATTG |
| HSI2_U12_A | CTGGAACGTCCACTCCCTAC |
