## Supplementary Figures for "*Arabidopsis TITAN LIKE* is required for U12-type intron splicing, especially of AT–AC subtypes"

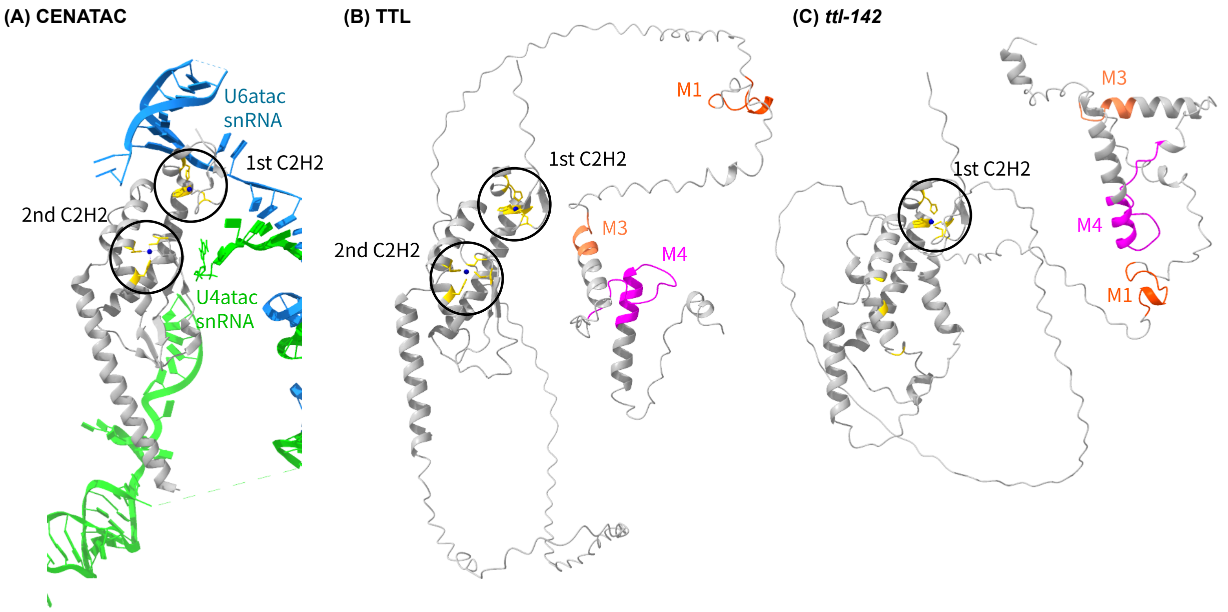


(D)

```
TTL        1   MKKPSKKSEIEFCTVCRFHHDQGSRHKYFPRHKSSLSSLLDRFRSKIADVRFFLKNPSVL   60
CENATAC    1   -MAPAQR-----CPLCRQTFFCGRGHVYSRKHQRQLKEALERLLPQVEAARKAIRAAQVE   54

TTL       61   RPQEQSQNRVWCVFCDEDIVELGSSFACSKA----INHFASSDHLKNIKQFLSKNGPAMD  116
CENATAC   55   RYVPEHERCCWCLCCGCEVREHLSHGNLTVLYGGLLEHLASPEHKKATNKFWWENKAEVQ  114

TTL      117   CIDEFRISEADVAKWEKKCQSFGNEDASFEGSCGQLSGTSNDIHTKLAFETMDRIKKVPA  176
CENATAC  115   MKEKFLVTPQDYARFKK---SMVKGLDSYE-----------EKEDKVIKEMAAQIREVEQ  160

TTL      177   HHINSYKSNDVMPLQYNTNEYQISLSEIPGVIHNGSYLNMDDSQFPLCDESGNGFGEHSI  236
CENATAC  161   SRQEVVRS-------------VLEPQAVP-----------DPEEGSSAPRSWKGMNSQVA  196

TTL      237   PCRSKDYSGNGNYCTQENYQVSQDKKQIDGSYNPPGVVGMTSISSSHSTDAGGNVHSGAP  296
CENATAC  197   SSLQQPSNLDLPPAPELDWMETGPSLTFIGHQDIPGV---------------GNIHSGAT  241
                                                                  M1
TTL      297   PPWLDANDGDFS-SVQLNQSDVARFQAKVPGKNRKLNPNRVGAAWAERRKIEIEMEKSGH  355
CENATAC  242   PPWMIQDEEYIAGNQEIGPSYEEFLKEKEKQKLKKLPPDRVGANFDHS------------  289
                               M2                    M3
TTL      356   VTKSNIDPDWLPNFGRVWQSGTRKESRKEFEKEKRKLVKTESISTESEPVKIQPYISKRA  415
CENATAC  290   ---SRTSAGWLPSFGRVWNNGRRWQSRHQFKTEAAAMKKQSHTEKS--------------  332
                        M4
TTL      416   RRESGE                                                        475
CENATAC  333   ------                                                        386
```

### Figure S1

Three-dimensional structural models of CENATAC and TTL. (A) Structure of CENATAC and two associated snRNAs extracted from the human minor spliceosome model (Protein Data Bank accession no. 8Y6O; https://www.rcsb.org/) (Bai et al. 2024). (B) Predicted structure of WT TTL generated using AlphaFold3 (Abramson et al. 2024). (C) Predicted structure of *ttl-142* according to AlphaFold3. The two C2H2 domains are circled, and histidine and cysteine residues are shown in yellow. Conserved motifs 1, 3, and 4 (M1, M3, and M4) are highlighted in orange, coral, and purple, respectively. (D) Alignment of TTL and CENATAC amino acid sequences. The deleted amino acid residues in *ttl-142* and *ttl-215* are underlined in red and coral, respectively.

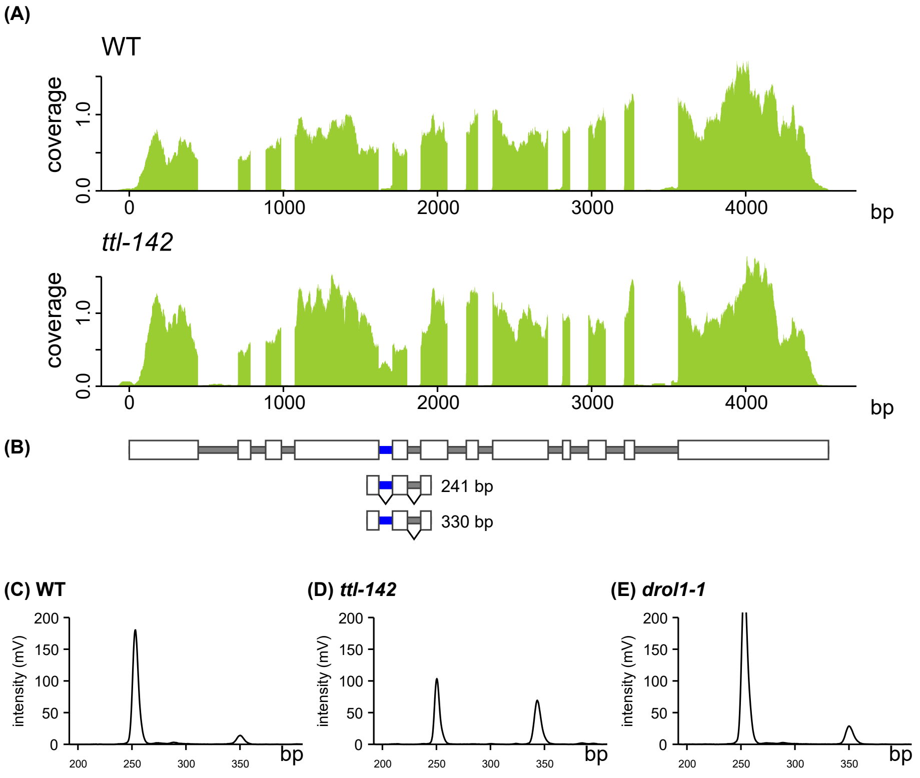


### Figure S2

RNA-Seq and RT-PCR analyses of retention of the U12-type GT–AG intron of HSI2 in WT and *ttl-142*. (A) RNA-Seq analysis of *HSI2*. All short reads were aligned to the *HSI2* gene sequence. The x-axis represents reference sequences, and the y-axis indicates short-read coverage. (B) Structure of the *HSI2* gene. Gray and white boxes represent introns and exons, respectively, and blue boxes indicate U12-type introns. Expected RT-PCR products shown in (C) to (E) are also indicated, together with their expected fragment sizes. (C–E) Microchip electrophoresis of DNA fragments amplified by RT-PCR from (C) WT, (D) *ttl-142*, and (E) *drol1-1* RNA. The x-axis represents DNA fragment size (bp), and the y-axis indicates fluorescence intensity corresponding to DNA abundance. See Figure 4 for details of the analytical procedures.

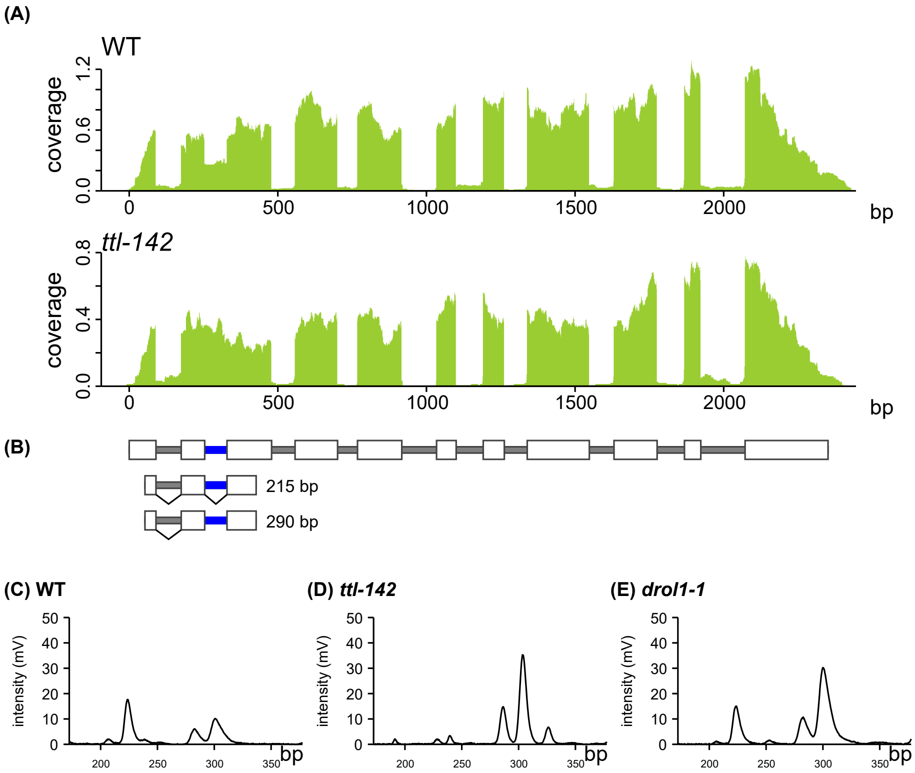


### Figure S3

RNA-Seq and RT-PCR analyses of retention of the U12-type GT–AG intron of *AT1G79880* (RNA recognition motif (RRM)-containing protein) in WT and *ttl-142*. (A) RNA-Seq analysis of *AT1G79880*. All short reads were aligned to the *AT1G79880* gene sequence. (B) Structure of the *AT1G79880* gene, with U12-type introns shown in blue. Expected RT-PCR products corresponding to (C) to (E) are indicated. (C–E) Microchip electrophoresis of DNA fragments amplified by RT-PCR from (C) WT, (D) *ttl-142*, and (E) *drol1-1* RNA. See Figure 4 for details of the analytical procedures.

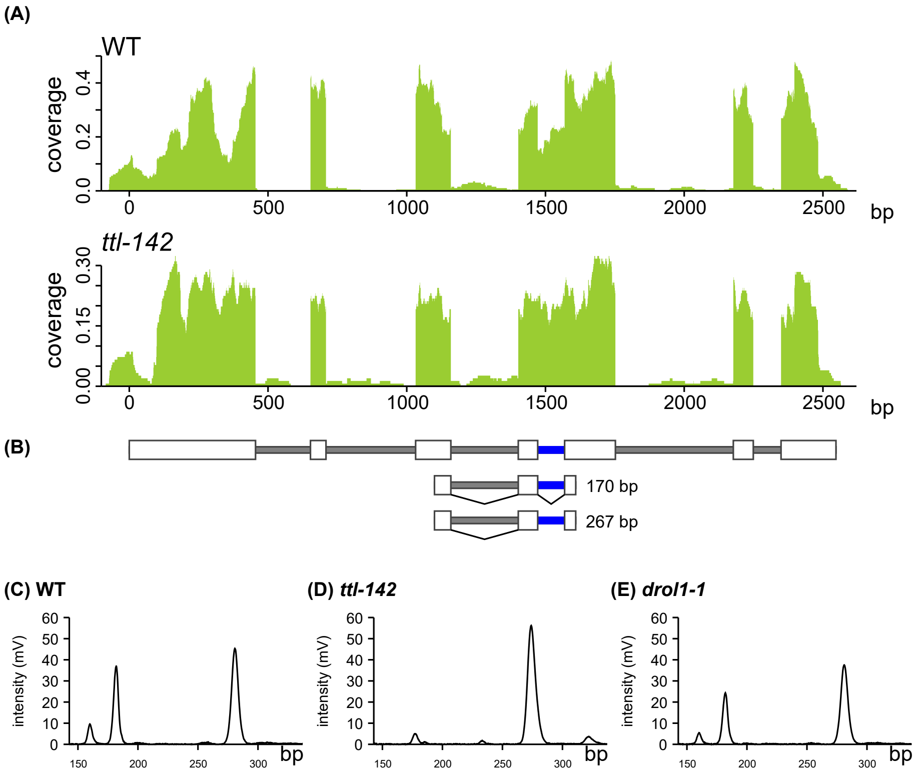


### Figure S4

RNA-Seq and RT-PCR analyses of retention of the U12-type GT–AG intron of *AT5G06620* (SET domain protein 38) in WT and *ttl-142*. (A) RNA-Seq analysis of *AT5G06620*. All short reads were aligned to the *AT5G06620* gene sequence. (B) Structure of the *AT5G06620* gene, with U12-type introns shown in blue. Expected RT-PCR products corresponding to (C) to (E) are indicated. (C–E) Microchip electrophoresis of DNA fragments amplified by RT-PCR from (C) WT, (D) *ttl-142*, and (E) *drol1-1* RNA. See Figure 4 for details of the analytical procedures.

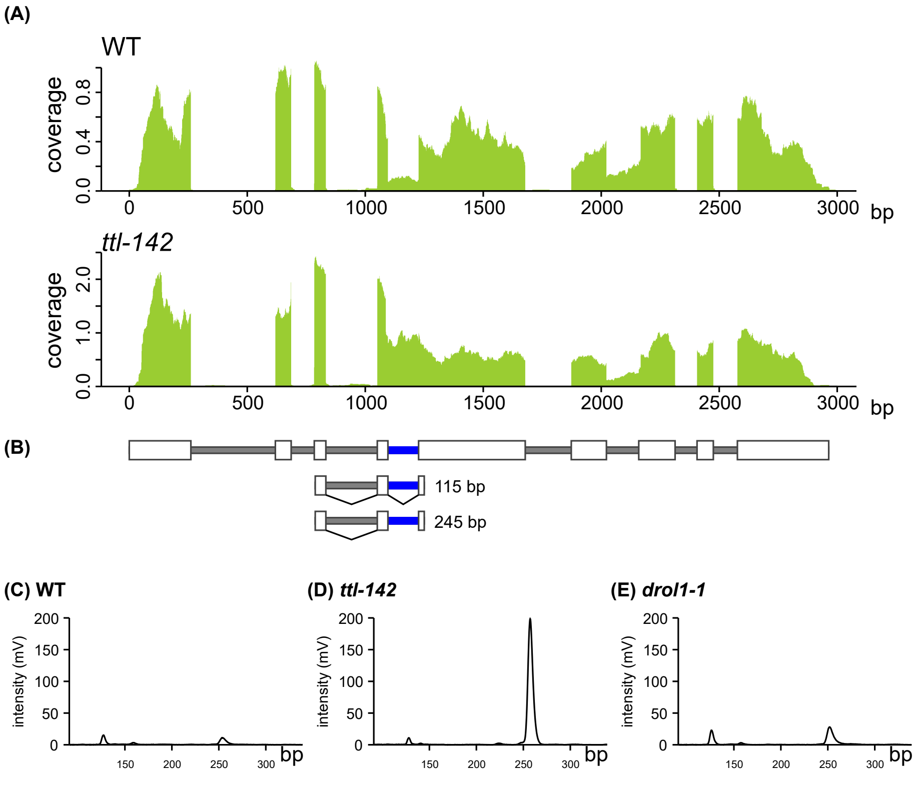


### Figure S5

RNA-Seq and RT-PCR analyses of retention of the U12-type GT–AG intron of *AT4G23330* (unknown protein) in WT and *ttl-142*. (A) RNA-Seq analysis of *AT4G23330*. All short reads were aligned to the *AT4G23330* gene sequence. (B) Structure of the *AT4G23330* gene, with U12-type introns shown in blue. Expected RT-PCR products corresponding to (C) to (E) are indicated. (C–E) Microchip electrophoresis of DNA fragments amplified by RT-PCR from (C) WT, (D) *ttl-142*, and (E) *drol1-1* RNA. See Figure 4 for details of the analytical procedures.

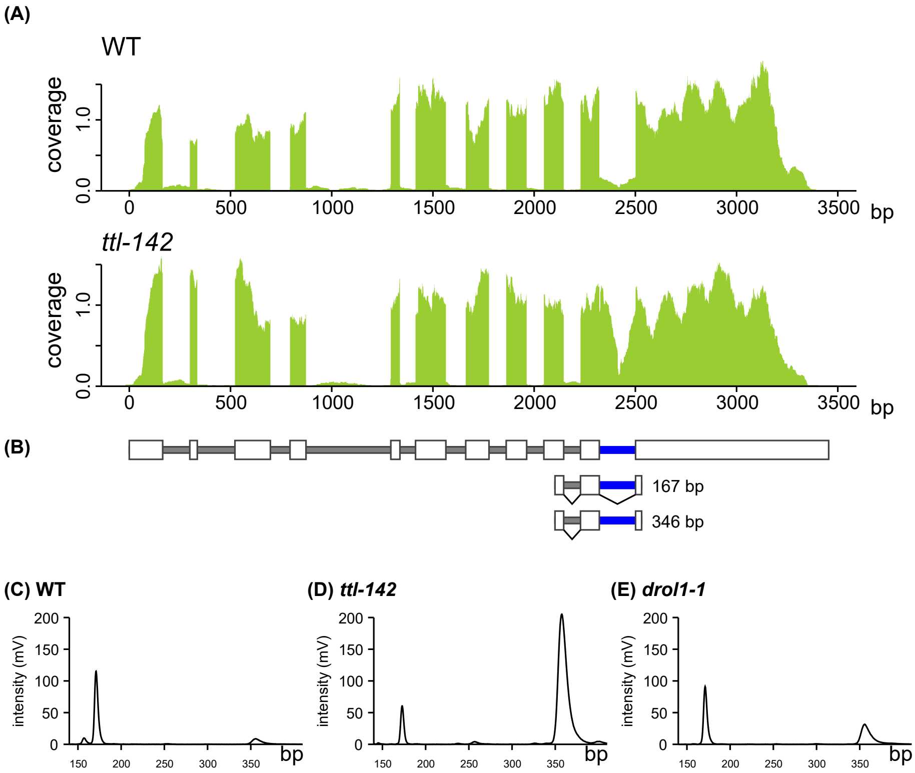


### Figure S6

RNA-Seq and RT-PCR analyses of retention of the U12-type GT–AG intron of *AT1G18090* (5′-3′ exonuclease family protein) in WT and *ttl-142*. (A) RNA-Seq analysis of *AT1G18090*. All short reads were aligned to the *AT1G18090* gene sequence. (B) Structure of the *AT1G18090* gene, with U12-type introns shown in blue. Expected RT-PCR products corresponding to (C) to (E) are indicated. (C–E) Microchip electrophoresis of DNA fragments amplified by RT-PCR from (C) WT, (D) *ttl-142*, and (E) *drol1-1* RNA. See Figure 4 for details of the analytical procedures.

```
TTL      480: CAGCTGTCTGGGACATCTAATGatatccacactaaacttgcgtttgaaactatggataga
TTLnmI   480: CAGCTATCTGGGACATCTAATGacattcacacgaagctggcgttcgagacgatggaccgg
         151:  Q  L  S  G  T  S  N  D  I  H  T  K  L  A  F  E  T  M  D  R

TTL      540: attaaaaaagttcctgcacaccatataaattcatataaatcaaatgatgttatgccttta
TTLnmI   540: atcaagaaggtgccggcccaccatataaattcgtataagtcgaatgatgtgatgccgttg
         171:  I  K  K  V  P  A  H  H  I  N  S  Y  K  S  N  D  V  M  P  L

TTL      600: cagtataatacGAATGAGTATCAGATATCACTTTCA
TTLnmI   600: cagtataatacGAATGAGTATCAGATATCACTTTCA
         191:  Q  Y  N  T  N  E  Y  Q  I  S  L  S
```

### Figure S7

Nucleotide sequence of TTLnmI. Alignment of the original TTL and TTLnmI nucleotide sequences surrounding the AT–AC intron with their corresponding deduced amino acid sequence. Exonic and intronic regions are indicated in uppercase and lowercase letters, respectively. All substituted nucleotides in TTLnmI are highlighted in yellow.
